# Life in the fast lane: Functional consequences of male-female dynamic differences in the renal auto-regulation of flow

**DOI:** 10.1101/2025.09.12.675896

**Authors:** Lingyun (Ivy) Xiong, Alan Garfinkel, Kevin M. Bennett, Edwin J. Baldelomar, Lauryn Brown, Kate Barrows, Volker Vallon, Aurelie Edwards, Alicia A. McDonough, Natalie Porat-Shliom, Eric J. Deeds

## Abstract

Tubuloglomerular feedback (TGF) is essential for the renal auto-regulation of flow. TGF is known to induce spontaneous oscillations in single-nephron tubular fluid flow in male kidneys. However, male-female differences in this dynamic behavior have not been studied. Leveraging intravital two-photon microscopy, resting-state magnetic resonance imaging, ultrasound-based and transdermal recordings, we found TGF-mediated oscillations across spatial scales in the rodent kidney, from single-nephron to whole-organ levels, and that male kidneys exhibited higher operating frequencies than females. To understand the mechanisms involved, we developed a dynamical systems model of TGF that agrees with physiological observations. Analysis of the mathematical model indicated that higher reabsorption rate and fluid flow efficiency in male proximal tubules not only result in higher frequencies, but also render male nephrons more susceptible to lose TGF-mediated oscillations. Furosemide abolished TGF-mediated oscillations in male kidneys and upregulated tubular injury marker KIM-1, suggesting that the propensity to lose TGF-mediated oscillations underlies the heightened risk for injury in males. Our analysis also indicated that SGLT-2 inhibition confers renoprotection by preserving TGF-mediated oscillations in hyperglycemia. Combining quantitative imaging and mathematical modeling, this study provides mechanistic insights into the transition from normal physiology to pathophysiology in the kidney.

## Introduction

According to the principles of homeostasis, physiological regulation maintains critical processes at steady states by negative feedback loops. However, a wide range of physiological systems exhibit oscillatory behaviors with well-defined frequencies, amplitudes and waveforms (1), including circadian rhythms (2, 3), core body temperature (4), plasma metabolite & hormone concentrations (5–9), the expression of transcription factors and kinases (10–16), as well as mitochondrial intermediates (17–19). Importantly, loss of oscillations has been shown to underlie the transition from health to disease: mutations in the p53 pathway promote cancer by abolishing p53’s oscillatory competence (20, 21); oncogenic Kras abrogates ERK oscillations to drive tissue hyperplasia (22); muscle-derived IL-6 pulses during exercise can be anti-tumor but prolonged release of IL-6 during inflammation is pro-tumor (23); exposure to chronic circadian disruption increases tumor burden (24); ultradian components of core body temperature rhythms are lost in aging and Huntington’s disease (25). Conversely, restoring rhythmic behavior can improve health: time-restricted eating improves cardiometabolic health in shift workers (26), and circadian alignment of caloric restriction promotes longevity in animal models (27). In this study, we investigate the role of physiological oscillations in renal function.

The kidney plays vital roles in maintaining a stable internal environment in the body. It not only removes waste, but also balances fluid volume and electrolyte composition (28). It does so by orchestrating filtration, fluid flow and solute reabsorption or secretion dynamically within each of its 1 million renal tubules (i.e., nephrons) in parallel. Receiving 20-25% of cardiac output, kidneys autoregulate renal blood flow (RBF) and glomerular filtration rate (GFR) over a wide range of renal perfusion pressures (80-180 mmHg), largely by tubuloglomerular feedback (TGF) and the myogenic response (29). TGF is important for maintaining GFR within a physiological range from minute to minute (30), and the myogenic response primarily protects glomerular capillaries against the damaging effect of increases in systolic blood pressure (31). Impaired renal auto-regulation has been identified in many pathophysiological conditions, including acute kidney injury, chronic kidney disease, diabetic kidney disease and hypertension (32–35).

As with the many examples of physiological oscillations described above, experimental measurements of tubular fluid flow and arterial blood flow in the rat kidney have repeatedly demonstrated robust oscillations (36–40), not steady states. The myogenic response and TGF modulate the vasomotor tone of the afferent arteriole (AA) in concert, but with different response times, enabling separation of signals in the time and frequency domain (41, 42). The faster myogenic response, occurring on the timescale of 5-10s (frequency: 0.1-0.2 Hz), adjusts the vascular tone of the AA to its intraluminal pressure via intrinsic mechano-transduction in the vascular smooth muscle cells (43). The slower process of TGF, occurring on the timescale of ∼30s (0.02-0.08 Hz), is triggered by changes in fluid flow and solute reabsorption along the proximal tubule (PT) and the thick ascending limb (TAL). Sodium chloride is sensed by macula densa (MD) cells at the end of the TAL, which in turn secrete positive (adenosine) or negative (prostaglandin E2 and nitric oxide) modulators to control vasoconstriction of the adjacent AA within the juxtaglomerular apparatus (44, 45).

One interesting aspect of renal function is the well-established notion that male and female kidneys exhibit distinct physiology, disease susceptibility and injury responses. The age-related decline of renal function is faster in men than in women (46–48). Age-matched male subjects (in both humans and rodents) have increased susceptibility to both acute kidney injury and chronic kidney disease (49–52). Comparison of transporter abundances along the renal tubule indicated a higher PT reabsorption capacity in male rodents than in females (53–56); especially relevant is the finding that Na/H exchanger 3 (NHE3), Na-phosphate cotransporter-2A (NaPi-2), and claudin-2 are more abundant and/or more active in the male PT. In contrast, distal tubule reabsorption capacity is higher in females than in males (56). Single-cell analysis of the mouse kidney has demonstrated that sexually dimorphic gene activity maps predominantly to PT segments (57–59), where male-biased lipid metabolism implies higher energy demands for reabsorption and increased reabsorption rate (58, 60). Sexually dimorphic gene expression has also been reported for the human kidney (58). Despite a recent report about salt-induced sex differences in TGF responsiveness at the MD (61), it is unclear how male and female kidneys differ dynamically in their auto-regulation of flow, which could be a key factor underlying the heightened risk for kidney diseases in males.

In this study, we examined male-female differences in the renal auto-regulation of flow across spatial scales, with a focus on TGF-mediated oscillations. By analyzing recordings from intravital imaging, resting-state MRI, ultrasound-based and transdermal measurements, we found that male kidneys exhibit higher frequencies than female kidneys, from the single-nephron level to the whole-organ level. To better understand how known molecular differences lead to these physiological phenomena, we developed a mathematical model of TGF that recapitulates nephron physiology. Analysis of the model provided a mechanistic explanation for male-female frequency differences, the heightened risk for tubular injury in males, as well as the renoprotective effect of SGLT-2 inhibition in diabetic kidney disease. This work highlights the physiological importance of oscillatory behaviors in the kidney, and demonstrates how a combination of experimental techniques and mathematical modeling can generate novel insights into the transition between normal physiology and pathophysiology.

## Results

### Physiological oscillations are observed across spatial scales in the male kidney

To systematically study TGF-mediated physiological oscillations *in vivo*, we employed multiple modalities to capture dynamic behaviors in the kidney at different spatial scales. For single-nephron tubular fluid flow, we used intravital two-photon (2P) microscopy (62–64) to track changes in fluorescence intensity of intravenously injected fluorescent probes within luminal regions of interest (ROIs) in early PT segments (**Figure 1A; Figure S1A**). Small-sized fluorophores (e.g., lucifer yellow) are often used to estimate average single-nephron GFR (SNGFR) (65, 66), but they are cleared by the kidney within minutes. In comparison, mid-sized fluorophores are suitable for hour-long tracking of tubular fluid flow (63), due to fractional glomerular filtration (67, 68). We found that luminal recordings with 70 kDa dextran-rhodamine in the kidney cortex exhibited robust oscillations in fluorescence intensity at the frequency of ∼0.03 Hz in 3-month-old male C57BL/6J mice (**Figure 1B**), which is associated with TGF-mediated oscillations and is independent of motion artifacts (**Figure S1A-B**). Considering that both glomerular filtration of solutes and tubular fluid flow are highly pulsatile, the interplay between these two processes likely contributes to the oscillations in fluorescent signals within a luminal ROI over time, allowing us to use relative changes in fluorescence intensity as a proxy for inferring TGF-mediated dynamics in SNGFR and tubular fluid flow (Methods). Detrended signals in adjacent luminal regions can also be used to estimate average tubular flow rate (**Figure S1C**), which is consistent with previous calculations in mice (65, 66). Hereafter, all intravital 2P imaging studies were performed with 70 kDa dextran-rhodamine.

**Figure 1.**
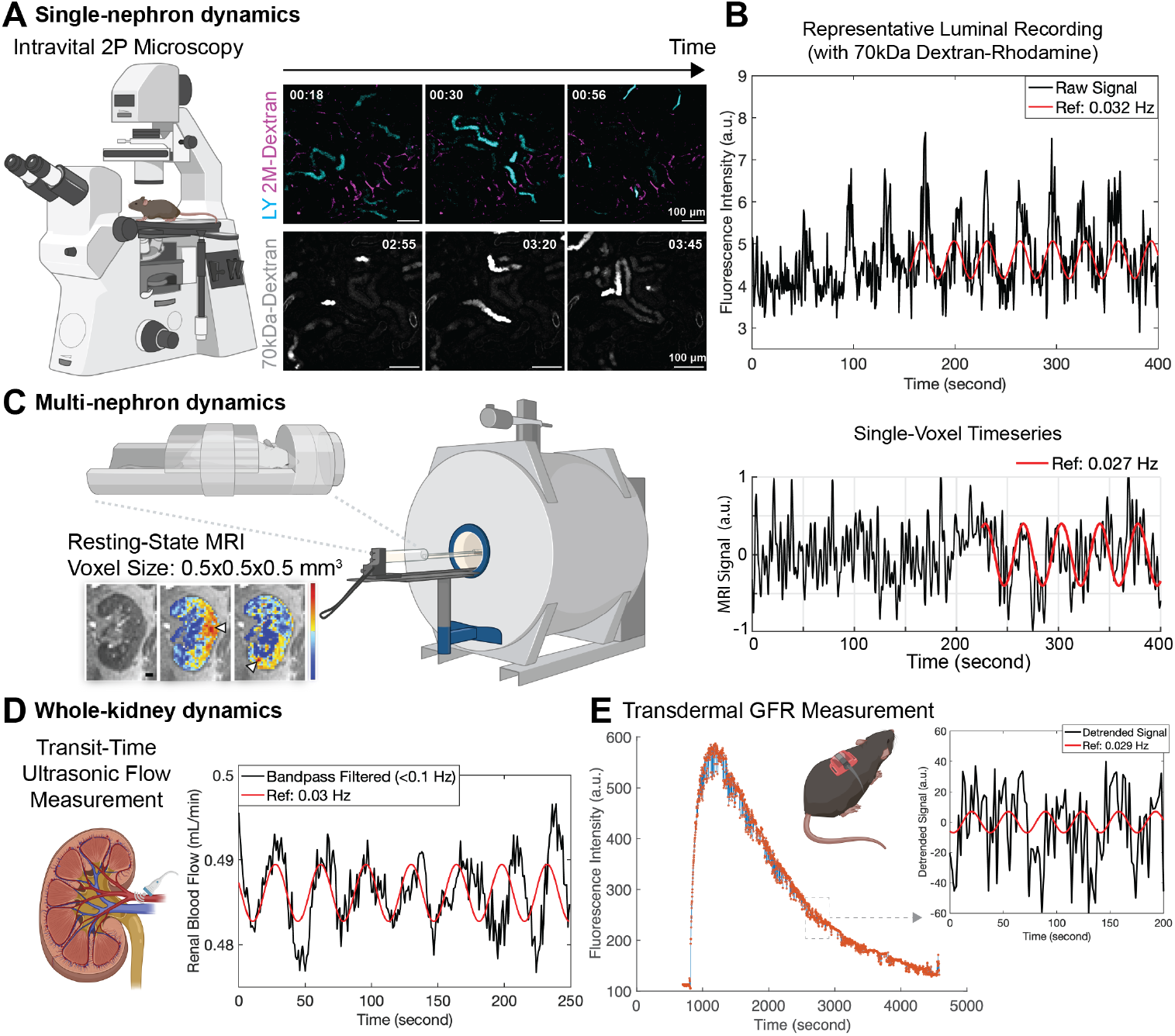
Physiological oscillations are observed across spatial scales in the male kidney. (A) Intravital two-photon (2P) microscopy was used to quantify single-nephron tubular fluid flow dynamics in the kidney cortex. Representative time-lapse images with Lucifer Yellow (LY) or 70kDa dextran-rhodamine show real-time changes in tubular fluid flow dynamics in the kidney of an anaesthetized 3-month-old male C57BL/6J mouse. 2M dextran-rhodamine was used together with LY to label the circulating plasma. (B) A representative recording of 70kDa dextran-rhodamine fluorescence intensity within a luminal ROI of the early PT segment in an anaesthetized 3-month-old male C57BL/6J mouse. Raw signals were overlaid with a reference sine wave of fixed frequency as indicated. (C) Resting-state magnetic resonance imaging (MRI) was used to detect physiological oscillations at the multi-nephron spatial scale. With a resolution of 0.5×0.5×0.5mm^3^ in a rat kidney, single-voxel recordings of MRI signals indicate collective dynamics among five to ten adjacent nephrons in the renal cortex (69). For illustration purposes, sagittal cross-section of a kidney and colormaps (left) were reprinted from ref. (*69*), with permission. The plot on the right represents a single-voxel timeseries obtained from a cortical region of an anaesthetized 3-month-old male Sprague-Dawley rat. Raw signals were overlaid with a reference sine wave. (D) Transit-time ultrasonic flow measurement was used to measure real-time renal blood flow through the renal artery. Data shown was collected from an anaesthetized 6-month-old male C57BL/6J mouse, previously reported in ref. (72). The original 1000Hz data was down-sampled to 2Hz and bandpass-filtered to visualize slow rhythms below 0.1 Hz, overlaid with a reference sine wave. (E) Glomerular filtration rate (GFR) can be estimated by fitting the transcutaneous clearance profile of a single intravenous bolus of FITC-sinistrin. Data shown was collected from an anaesthetized 3-month-old male C57BL/6J mouse, previously reported in ref. (76). The raw data points, with a 0.5 Hz sampling rate, are colored in red. Detrended signals for a 200-second interval are shown on the right, overlaid with a reference sine wave.

To examine TGF-mediated oscillations at larger spatial scales, we used resting-state magnetic resonance imaging (MRI) to detect physiological oscillations in the rat kidney. We found reproducible oscillatory MRI signals at ∼0.03 Hz across the kidney cortex in 3-month-old male Sprague-Dawley (SD) rats (**Figure 1C**). With a spatial resolution of 0.5×0.5×0.5mm^3^, periodic single-voxel MRI signals indicate collective oscillatory changes among 5-10 neighboring nephrons (69), suggesting regional synchronization of activities related to fluid flow and oxygenation, as has been reported for blood flow in star vessels across the kidney surface (70, 71). Next, we re-analyzed published ultrasound-based measurements of blood flow through the renal artery (72), and detected oscillations in renal blood flow at ∼0.03 Hz in 6-month-old male C57BL/6 mice (**Figure 1D**), suggesting that TGF-mediated oscillations can be detected at the whole-organ level (73–75). Moreover, published recordings from transdermal GFR measurement experiments in 3-month-old male C57BL/6J mice (76), tracking the whole-body clearance profile of FITC-sinistrin (77), often display oscillatory fluorescence signals at ∼0.03 Hz during the exponential decay phase (**Figure 1E**), which can be attributed to oscillatory GFR (**Figure S1D**). Together, these *in vivo* experimental measurements demonstrated physiological oscillations at ∼0.03 Hz across spatial scales in the male kidney.

### Male-female frequency differences in TGF-mediated oscillations

Using intravital 2P microscopy, we compared single-nephron dynamics in tubular fluid flow between male and female C57BL/6J mice of different age groups. Two representative recordings from 3-month-old mice are shown in **Figure 2A**: fluorescence signals in a male early PT segment showed an oscillatory pattern with a dominant frequency of 0.043 Hz, while signals in a female early PT segment showed a dominant frequency of 0.025 Hz. Surveying multiple early PT regions in each animal, we noted larger variations in the frequency distribution among male tubules (spanning 0.03-0.08 Hz) than in females (0.015-0.04 Hz) (**Figure 2B**; **Figure S2A**), with male tubules having a significantly higher median frequency (0.046 Hz) than females (0.023 Hz) (*P* < 10^−6^, Methods). Similar frequency differences were observed in 15-month-old mice, but not in 1-month-old mice (**Figure 2B**). Previously, we showed that sexually dimorphic gene expression in the mouse kidney manifests between 1 and 2 months of age, which was driven by testosterone and the androgen receptor (58). Our findings suggest that gonadal hormones may also play an important role in determining the frequency of TGF-mediated oscillations in tubular fluid flow.

**Figure 2.**
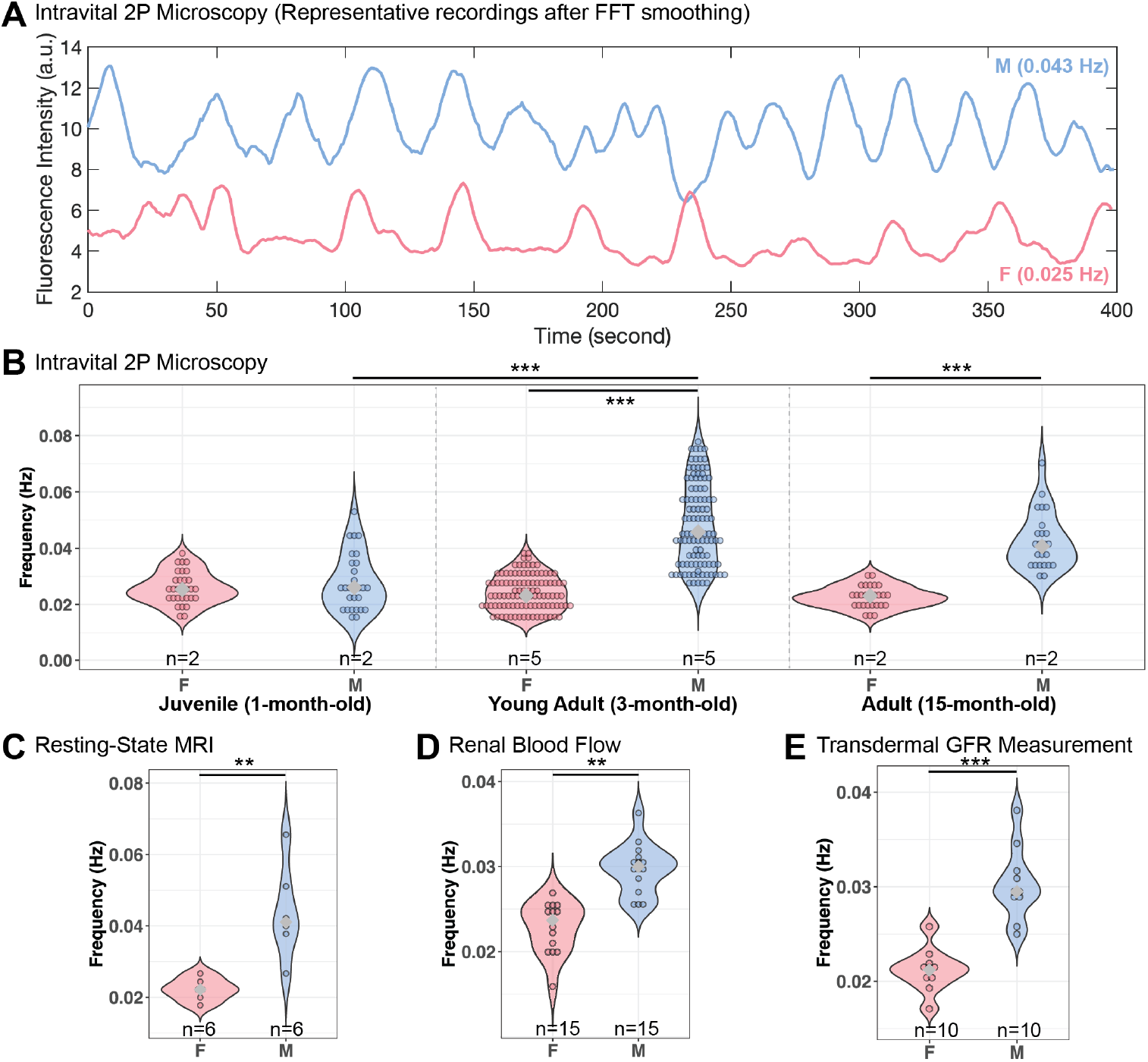
Male-female frequency differences in TGF-mediated oscillations. (A) Representative recordings of intravital 2P microscopy showing Fast Fourier Transform (FFT)-smoothed signals of 70kDa dextran-rhodamine fluorescence over time within early PT segments in 3-month-old male and female C57BL/6J mice. Dominant frequencies are indicated in parentheses. (B) Distributions of single-nephron dominant frequencies in male and female C57BL/6J mice of three age groups (1-, 3- and 15-month-old), measured by intravital 2P microscopy. Dominant frequencies in each tracing were determined by FFT. Over a dozen regions were quantified in each animal; individual values for each region are shown. (C) Distribution of multi-nephron dominant frequencies in 3-month-old male and female SD rats, measured by resting-state MRI. Dominant frequencies in each single-voxel timeseries were determined by FFT. In each animal, 250-300 cortical voxels were quantified, and the mode (most likely value) of the dominant frequency distribution is reported. (D) Distribution of whole-kidney dominant frequency in 6-month-old male and female rodents (C57BL/6 and C57BLKS/J mice; Wistar Kyoto rats), quantified from transit-time ultrasound-based renal blood flow data previously reported in ref. (72, 78). Dominant frequencies in each recording were determined by FFT. (E) Distribution of whole-body clearance dominant frequency in male and female C57BL/6J mice (3- and 7-month-old), quantified from transdermal GFR measurement previously reported in ref. (76, 79). Dominant frequencies in each recording were determined by FFT. In all cases, sample number *n* indicates total numbers of animals used. Statistical significance was determined by resampling (Methods; * *P* < 0.05; ** *P* < 0.01; *** *P* < 0.001).

Going beyond single nephrons, we also observed similar male-female frequency differences at the multi-nephron and whole-kidney levels. Surveying a large number of voxels in the kidney of 3-month-old male and female SD rats, resting-state MRI detected oscillatory signals across cortical regions in both sexes. We found that male rats have a higher median frequency (0.041 Hz) than in females (0.022 Hz) (*P* = 0.0063, Methods; **Figure 2C**). Next, by re-analyzing renal blood flow recordings in 6-month-old male and female C57BL/6 mice and Wistar Kyoto rats (72, 78), we found that male animals consistently show a higher median frequency (0.030 Hz) than females (0.022 Hz) within the TGF frequency range (*P* = 0.0026, Methods; **Figure 2D**; **Figure S2B**). Lastly, after extracting dominant frequencies from recordings during transdermal GFR measurements in 3- and 7-month-old male and female C57BL/6J mice (76, 79), we also found a higher median frequency in males (0.030 Hz) than in females (0.021 Hz) (*P* = 6.7x 10^−5^, Methods**; Figure 2E**; **Figure S2C-D**). Across spatial scales, we identified male-female frequency differences in TGF-mediated oscillations in the kidney, demonstrating that in normal conditions, male kidneys operate “in the fast lane” while maintaining auto-regulation of flow.

### A compartmental model of TGF recapitulates essential nephron physiology

To investigate the mechanisms underlying these male-female frequency differences, we developed a dynamical systems model of TGF (Methods). Dividing the PT and TAL into arrays of smaller units, our compartmental model of TGF keeps track of fluid volume in individual PT compartments and sodium concentration in individual TAL compartments over time, in the form of linked Ordinary Differential Equations (**Figure 3A**). In particular, we stipulate two parameters governing tubular fluid flow and sodium handling in the PT: a fluid flow rate constant *κ*, which determines flow efficiency across compartments, and a sodium reabsorption rate *λ* (see Methods), featuring flow-dependent reabsorption (80–82). The negative feedback from the MD to the AA controls the rate of fluid flow feeding into the first PT compartment, thus closing the negative feedback loop in the tubule (**Figure 3A**). Conceptually, this model encapsulates the negative feedback between GFR and sodium concentration sensed by the MD with an intrinsic time delay, only that the delay is not imposed explicitly by the modeler, but is implicit, inherent in the fact that fluid takes time to travel from the glomerulus to the MD. The sensitivity of TGF is modeled by the steepness of the sigmoidal function that relates sodium chloride concentration sensed by the MD to changes in SNGFR (**Figure 3A-B**). When parameterized to the specifications of mouse nephrons (see Methods), model simulations reproduce self-sustained sawtooth-like oscillatory behavior in SNGFR over time, as observed experimentally (**Figure 1B; Figure S3A**). It has been shown that TGF sensitivity at the MD plays an important role in determining the oscillatory state of the system (83–85). The compartmental model also exhibits this feature: switching from steep feedback to shallow feedback converted oscillatory SNGFR to constant SNGFR (**Figure 3B**).

**Figure 3.**
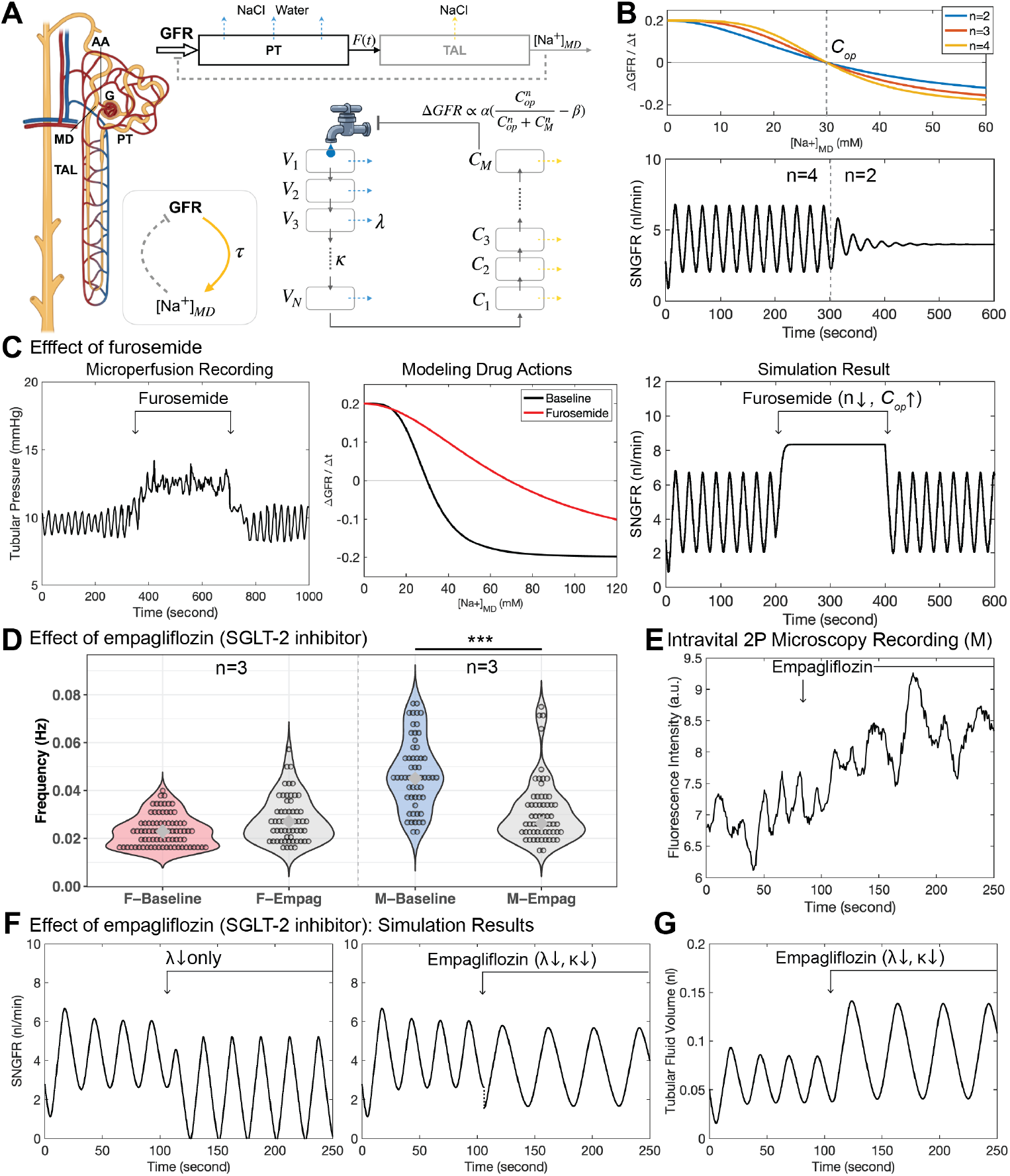
A compartmental model of TGF recapitulates essential nephron physiology. (A) Schematic of the compartmental model of TGF (see Methods). (B) TGF sensitivity at the macula densa (MD) is modeled by the maximal slope of the sigmoidal function (top), which plays a major role in determining system behavior (bottom): a ‘steep’ slope (n=4) sustains oscillatory SNGFR, while a ‘shallow’ slope (n=2) abolishes oscillations and sustains constant SNGFR. (C) Dynamic responses in proximal intra-tubular pressure during furosemide infusion (left; plot was redrawn from ref. (87), with permission) can be replicated by adjusting model parameters informed by the known drug action that furosemide inhibits NKCC2 to block sodium sensing at the MD (middle). Simulation result produced by modifying the sigmoidal function for the indicated interval is shown (right). (D) Acute effect of empagliflozin (50 *μ*g/kg; intravenous bolus injection) on single-nephron dominant frequency in 3-month-old male and female C57BL/6J mice. Recordings were obtained using intravital 2P microscopy, and dominant frequencies of tubular fluid flow dynamics in early PT segments were determined by FFT. Statistical significance was determined by resampling (Methods; *** *P* < 0.001). (E) A representative recording of intravital 2P microscopy shows that empagliflozin injection triggered an elevation in fluorescence intensity and a decrease in frequency in a 3-month-old male C57BL/6J mouse. (F) Simulated SNGFR over time with lowering *λ* by 40% alone (left) or simultaneously lowering *λ* by 40% and lowering *κ* by 50% (right). (G) Simulated tubular fluid volume in an early PT segment over time with simultaneously lowering *λ* by 40% and lowering *κ* by 50%.

We tested the validity of the compartmental model by simulating the effect of the loop diuretic furosemide. Experimentally, Yip et al. showed that 0.2 mM furosemide perfused into a male rat nephron abolished TGF-mediated oscillations in intratubular pressure, and induced constant elevated pressure (86, 87) (**Figure 3C**, left). Furosemide is known to block sodium chloride sensing in MD cells by inhibiting the luminal sodium potassium chloride cotransporter (NKCC2) (88). This action of furosemide can be modeled by increasing the operating concentration and switching from steep to shallow feedback at the MD (**Figure 3C**, middle), yielding a simulation that agrees with the experimental recording (**Figure 3C**, right). By modulating drug action at the MD, the compartmental model can also reproduce the partial-inhibition effect of a lower dose of furosemide (**Figure S3A-C**).

Like furosemide, SGLT-2 inhibitors have also been shown to influence TGF in modulating renal auto-regulation of flow. To study how SGLT-2 inhibition (SGLT-2i) impacts TGF-mediated oscillations, we injected the SGLT-2 inhibitor empagliflozin (50 *μ*g/kg) intravenously and compared tubular fluid flow dynamics before and after treatment, using intravital 2P microscopy. We found that empagliflozin lowered the frequency of TGF-mediated oscillations in 3-month-old male C57BL/6J mice within minutes (*P* < 10^−6^, Methods; **Figure 3D**). Higher mean fluorescence intensities implied increased flow, due to osmotic diuresis (89), accompanied by gradual slowing down in rhythmic behavior (**Figure 3E**). While SGLT-2 inhibitors are known to inhibit sodium glucose co-transport in the PT (90, 91), analysis of our mathematical model revealed that decreasing proximal sodium reabsorption rate (*λ*) alone was insufficient to produce frequency change (**Figure 3F**, left). In contrast, a concurrent decrease in both *λ* and *κ* is required to recapitulate the experimental observation (**Figure 3F-G**). This finding suggests that an increase in the impedance to flow of the luminal fluid (due to glucose accumulation and increased viscosity and osmolarity) is crucial for inducing frequency changes in tubular fluid flow. Interestingly, acute treatment of empagliflozin in 3-month-old female C57BL/6J mice did not result in significant changes in frequency (**Figure 3D, S3D**).

### Loss of TGF-mediated oscillations underlies heightened risk in males

To investigate why male and female nephrons show different frequency distributions in TGF-mediated oscillations, we studied how the behaviors of the compartmental model depend on key parameters. Considering that male and female PTs differ in their sodium transport and metabolic capacities (56, 58), we reasoned that proximal sodium reabsorption rate (*λ*) may be an important parameter for determining system behavior. We have also seen that the fluid flow rate constant (*κ*) modulates the frequency of tubular fluid flow. We thus characterized the behavior of the model across a wide range of *λ*-*κ* combinations.

The distribution of average SNGFR covers the full range across the *λ*-*κ* space: low-*λ* and high-*κ* maintain low flow, while high-*λ* and low-*κ* maintain high flow (**Figure 4A**). Our simulations indicate that oscillatory behaviors are sustained for a subset of *λ*-*κ* combinations showing moderate average SNGFR, where higher *λ* generally results in diminishing amplitude (**Figure 4B**). As in the case of simulating the effect of SGLT-2i, the value of *κ* largely determines the frequency (**Figure 4C**), where greater *κ* corresponds to higher frequency. Male PTs have larger lumen size than females (92), suggesting that male tubules tend to have higher *κ* (Methods). Since male and female PTs have a fractional reabsorption of 0.65-0.8 and 0.5-0.6, respectively (53, 93), we can map the regions in *λ*-*κ* space where male and female tubules likely operate. Matching the median frequency in male and female kidneys (**Figure 2B**), region (c) in **Figure 4C** represents a typical *λ*-*κ* combination for male tubules, and region (e) for females. Corresponding simulations of SNGFR are shown in **Figure 4D**, in which male parameter values (**Figure 4D-c**) are ∼2-fold (for *κ*) and 3-fold (for *λ*) of female parameter values (**Figure 4D-e**). Previous research showed that administration of benzolamide or acetazolamide in male animals acutely reduced average SNGFR by directly decreasing *λ* (94, 95), thus transitioning from region (c) to (b), where average SNGFR is lowered without changes in frequency (**Figure 4C**). Further decreases in *λ* would result in constant low flow [e.g., region (a)], which could be the case in ischemia-reperfusion injury (76), or in the case of claudin-2 deficiency and associated kidney stone disease (96).

**Figure 4.**
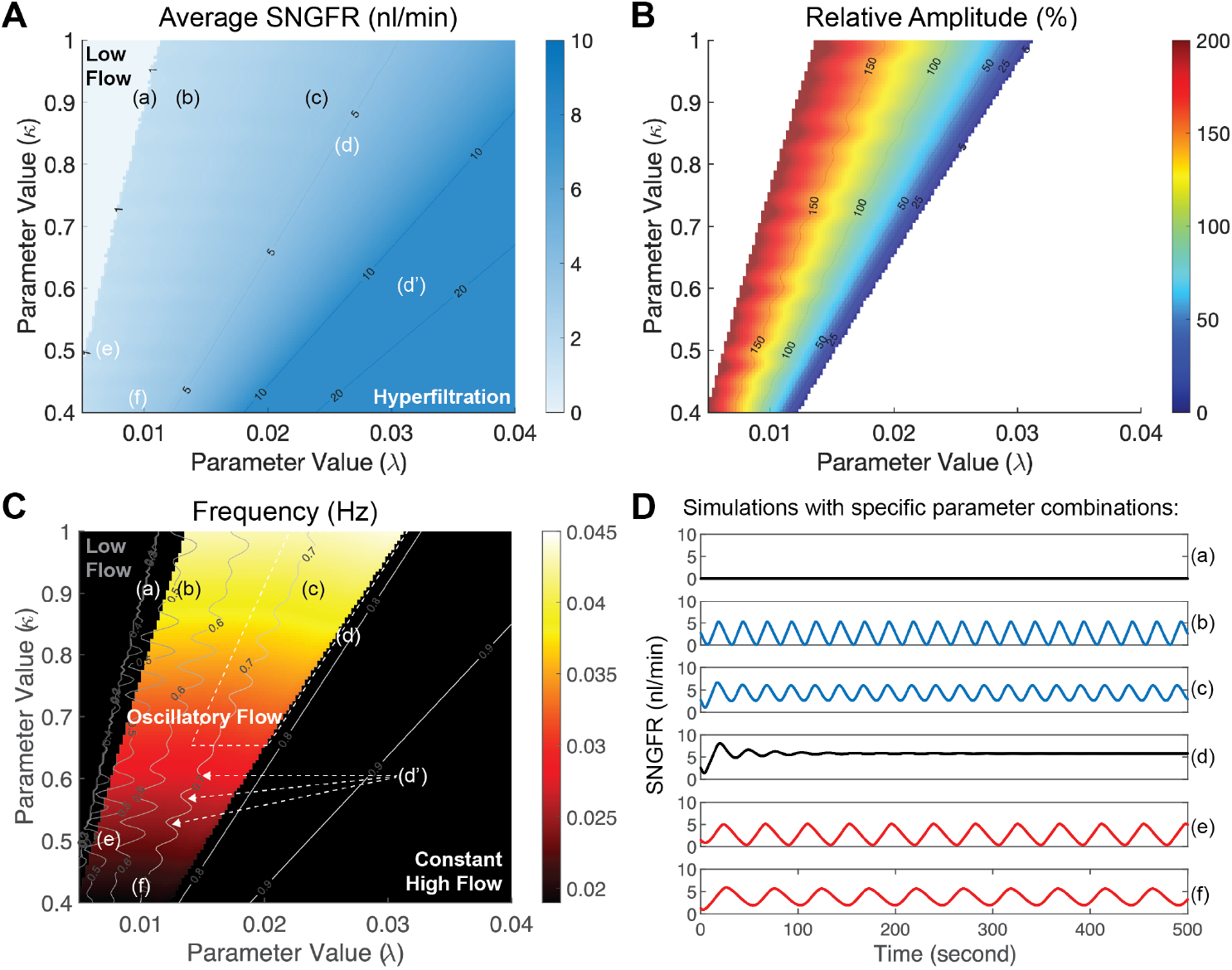
Loss of TGF-mediated oscillations underlies heightened risk in males. (A) Numerical simulations were performed for different combinations of *λ* and *κ*, while keeping a ‘medium’ slope (n=3) of the sigmoidal function at the MD. The heatmap shows the distribution of average SNGFR across the 2D parameter space for specific *λ*-*κ* combinations. (B) Heatmap indicates the amplitude of oscillatory behavior relative to the average SNGFR, revealing a region in *λ*-*κ* space where TGF-mediated oscillations are self-sustained. (C) Heatmap indicates the frequency of oscillatory behavior as shown in (B), overlaid with the contour plot indicating fractional reabsorption of sodium in the PT. (D) Simulation results with specific parameter combinations as marked in panel (A) and (C). For a given value of *κ, λ* needs to be sufficiently large to induce TGF-mediated oscillations [contrasting (a) and (b)], but high *λ* risks losing TGF-mediated oscillations. Parameter combination (c) represents a typical nephron with male frequency & PT fractional reabsorption, while combination (e) represents a typical nephron with female frequency & PT fractional reabsorption. Mild perturbation in *λ* and *κ* (such as in the case of prolonged high blood glucose) can result in loss of TGF-mediated oscillations in male nephrons [e.g., transition from (c) to (d)] but not in female nephrons [e.g., transition from (e) to (f)], suggesting that male nephrons are more prone to lose TGF-mediated SNGFR oscillations. Parameter combination (d’) represents a typical male nephron in type 1 diabetes, estimated from reported measurements of average SNGFR in *Ins2*^*+/Akita*^ mice (66). Parameter combinations: (a) *λ* = 0.01, *κ* = 0.92 ; (b) *λ* = 0.013, *κ* = 0.92; (c) *λ* = 0.024, *κ* = 0.92; (d) *λ* = 0.026, *κ* = 0.86; (e) *λ* = 0.008, *κ* = 0.5; (f) *λ* = 0.01, *κ* = 0.44 [units: 10 *s*^−1^].

Note that male tubules operate close to the right-hand boundary of the oscillatory regime, where a slight change in *λ* or *κ* can take the system out of self-sustained oscillations and into a state of constant high flow, in which TGF-mediated auto-regulation of flow is attenuated. For instance, prolonged high blood glucose could result in increased *λ* (upregulation of SGLT-2) (97) and decreased *κ* (osmotic diuresis), causing a loss of the capacity to maintain oscillatory behaviors [e.g., a transition from region (c) to (d)], as reported for short exposure to hyperglycemia in male SD rats (69). On the other hand, the same shift in *λ*-*κ* space from region (e) to (f) keeps female tubules operating within the oscillatory regime, suggesting that female tubules are protected from hyperglycemia-induced loss of TGF-mediated oscillations.

SNGFR in male *Ins2*^*+/Akita*^ mice was shown to be significantly higher than those in C57BL/6 controls (66), suggesting that male tubules in type 1 diabetes operate in the region of high-*λ* and low-*κ* [e.g., region (d’) in **Figure 4C**], risking hyperfiltration, metabolic stress and glomerular damage (33, 98, 99). In this condition, SGLT-2i can confer renal protection by bringing the system back into or keep the system in the oscillatory region and maintain moderate SNGFR (66). A recent study showed that luminal glucose can activate SGLT-1 in MD cells, triggering NOS1-dependent nitric oxide production and lowering TGF responsiveness in male mice (100). We simulated this scenario with shallower feedback in the compartmental model, to find that the oscillatory regime in the *λ*-*κ* phase diagram is greatly reduced in width and left-shifted (**Figure S4A**). This indicates that male tubules are even more vulnerable to lose TGF-mediated oscillations during hyperglycemia, resulting in increased risk for hyperfiltration and associated damage. TGF sensitivity at the MD is also blunted in the aging male kidney (34), where a narrowing of the oscillatory regime indicates a case of what has been described as “homeostenosis” (101) that potentially underlies aging-related decline in renal function. In contrast, female tubules remain in the oscillatory regime in spite of shallower feedback (**Figure S4A**), implying that female kidneys can tolerate a wider range of TGF sensitivity to achieve dynamic auto-regulation.

### Acute loss of TGF-mediated oscillations upregulates tubular injury marker

To test whether acute loss of TGF-mediated oscillations has any physiological consequences, we treated 3-month-old male and female C57BL/6J mice with furosemide (1 mg/kg; intravenous bolus injection) during intravital 2P microscopy. We found that furosemide triggered a complete inhibition of TGF-mediated oscillations in male mice (**Figure 5A**). 30 minutes post furosemide injection, we collected kidneys for fixation and immunostaining. Male kidneys treated with furosemide showed increased expression of the tubular injury marker KIM-1 in PT segments (F-actin stains the brush boarder) when compared to controls (*P* < 10^−6^, Methods; **Figure 5C-D**), indicating that the injury occurred upstream of NKCC2 inhibition in the TAL. The extent of KIM-1 upregulation is comparable to the effect of 30-minute unilateral ischemia-reperfusion injury in male mice (102). Low-dose furosemide infusion in male rats was reported to induce tubular necrosis within 2 hours (103), indicating that loss of TGF-mediated oscillations indeed co-occurred with tubular injury.

**Figure 5.**
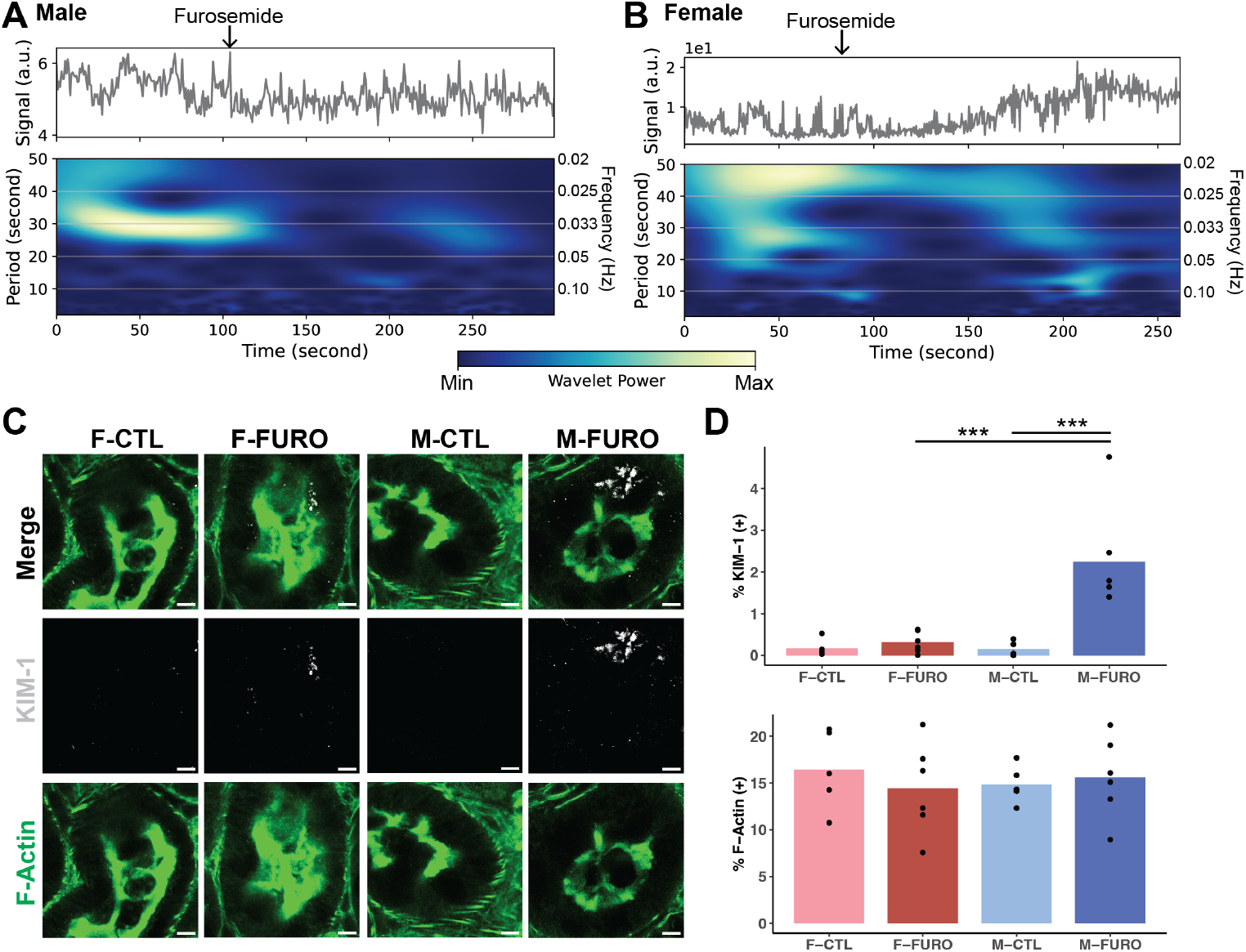
Acute loss of TGF-mediated oscillations upregulates tubular injury marker. (A-B) Representative recordings from intravital 2P microscopy demonstrate the effect of furosemide (1 mg/kg; intravenous bolus injection) on tubular fluid flow dynamics in 3-month-old male (A) and female (B) C57BL/6J mice. Wavelet analysis indicates the time-frequency components over time in each tracing. (C) Male and female C57BL/6J mice were treated with intravenous injection of furosemide (FURO; 1 mg/kg) or vehicle (CTL) for 30 minutes. Fixed kidney sections were stained for KIM-1 (grey) and F-actin (green) and imaged at 63X magnification. Scale bar, 5 *μ*m. (D) Quantification of stain-positive areas (n = 5-6) in each condition is shown for KIM-1 and F-actin. Statistical significance was determined using resampling (Methods; *** *P* < 0.001).

In contrast, female kidneys showed little inhibition of TGF-mediated oscillations after furosemide treatment, in spite of an elevation in fluorescence intensity (**Figure 5B**). Simulations with our compartmental model also indicated that female tubules are more resilient to furosemide-induced loss of TGF-mediated oscillations (**Figure S3C**). KIM-1 expression in furosemide-treated female kidneys did not show significant increases compared to controls (**Figure 5C-D**). These findings suggest that female kidneys are protected from loss of TGF-mediated oscillations and associated tubular injury, resonating with earlier reports that female kidneys are more resilient to acute kidney injury (104, 105).

## Discussion

While physiological oscillations have been observed in many systems, it has been a challenge to interpret these dynamic behaviors, both in terms of their mechanisms and their functional roles. In the male rat kidney, physiological oscillations at ∼0.03 Hz have long been noted in tubular fluid flow and arterial blood flow (36–40). These oscillations have been attributed to the action of TGF, a negative feedback loop that autoregulates renal blood flow to minimize the impact of blood pressure fluctuations (35). Here we show that physiological oscillations at frequencies associated with TGF are detected across a wide range of spatial scales, from the single nephron imaged by intravital microscopy, to those measured by clinically-relevant radiological imaging and transdermal GFR estimation. We show that male-female differences in the frequencies of these oscillations are common across all measurement modalities. Finally, we show that furosemide administration abolishes TGF-mediated oscillations and lead to heightened risk of kidney injury in males, likely due to sex differences in how furosemide affects TGF-mediated dynamics. To understand these findings in the context of nephron physiology, we developed a testable mathematical model of TGF-mediated oscillations, laying the groundwork for studies aimed at facilitating individualized monitoring of chronic kidney disease progression and responses to therapies.

### Towards an understanding of heightened risk in males

Previous studies of TGF-mediated oscillations predominantly focused on male animals; here we extended our study to examine both male and female animals. We found significant male-female frequency differences in TGF-mediated oscillations across spatial scales, which cannot be inferred from molecular data alone. Based on our mathematical model of TGF, we found that the higher frequency in male tubules may result from a combination of higher proximal sodium reabsorption rates and tubular fluid transport efficiencies (i.e., lower impedances to tubular fluid flow). Analysis of the model revealed that compared to females, male tubules operate close to the boundary of the oscillatory regime, which, upon perturbation such as exposure to hyperglycemia, can transition to the state of constant high flow, risking hyperfiltration and tubular injury. In contrast, female tubules are protected from loss of TGF-mediated oscillations, in part explaining why females are more resilient to injury. To better understand female protection from kidney diseases, additional studies are warranted to characterize TGF-mediated oscillations in female kidneys during pharmacological interventions and in disease models.

One might argue that we have not shown that the loss of oscillatory activity, for example, by furosemide administration, is the *cause* of renal injury, only that they co-occurred. But the same co-occurrence is seen in the loss of oscillation (and subsequent kidney injury) in other pathophysiological conditions, such as the Spontaneously Hypertensive Rat, in which loss of TGF-mediated oscillation and low-flow conditions are accompanied by kidney damage (37, 106). It is plausible that the final common pathway uniting these disparate mechanisms is the loss of oscillation. Our findings relate to reports that male kidneys are more prone to acute kidney injury (105, 107), oxidative stress and ferroptosis in the renal tubule (108–110), hypertensive kidney injury (111) and clear cell renal cell carcinoma (112–114), as well as worse progression in polycystic kidney disease (115–117). Elucidating the role of TGF-mediated oscillations in these pathophysiological transitions would help uncover an under-appreciated dimension of disease mechanisms.

### SGLT-2i and renal protection

SGLT-2i is recommended as the frontline treatment for diabetic kidney disease (118–120). Recently, SGLT-2i has also proved, in a clinical trial, to slow the decline of physical health in patients with chronic kidney disease (121). Previous pre-clinical studies predominantly investigated the effect of SGLT-2i on steady-state GFR in diabetic animal models, but questions regarding how SGLT-2i impacts TGF-mediated oscillations, or whether male and female kidneys respond differently, remain under-studied. Acute injection of an SGLT-2 inhibitor significantly lowered the frequency of TGF-mediated oscillations in male mice, which was replicated by decreasing both the proximal sodium reabsorption rate and the rate constant of tubular fluid flow in our compartmental model of TGF. Mathematical analysis demonstrated that the renal protective effect of SGLT-2i in diabetic kidney disease not only involves lowering average SNGFR, but could also involve restoring TGF-mediated oscillation and preventing the loss of oscillatory behavior. Besides providing mechanistic insights into the observed male-female frequency differences, our compartmental model of TGF also represents a computational tool for the *in-silico* investigation of drug actions for personalized medicine.

Studies of drugs, including SGLT2i, have revealed that patient responses to therapies are highly variable (120). There is a strong need to monitor responses to these therapies in individual patients, and to identify optimal therapeutic strategies for preventing chronic kidney disease. Because the modulation of TGF is a key protective mechanism of SGLT2i (33), the application of new, noninvasive tools to ascertain TGF responses could provide valuable information about an individual’s disease progression and therapy response. A recent pilot study demonstrated that it is feasible to use resting-state MRI for detecting TGF-mediated oscillations in human kidneys (69). Thus it is increasingly possible to perform timely and accurate assessment of renal function in humans and evaluate real-time patient response to SGLT-2 inhibitors and other medications (122, 123). Furthermore, transdermal GFR measurement has proven to be an accurate method for point-of-care assessment of real-time GFR in humans (124), opening a possibility to incorporate GFR dynamics into standard-of-care practices.

### Functions of TGF-mediated oscillations

One of the obstacles to a wider awareness of oscillatory physiology has been a lack of understanding of the physiological role that these oscillations play (125). In the kidney, oscillations in the partial pressure of oxygen occur at the same frequency as TGF-mediated oscillations in tubular fluid flow (126), suggesting that TGF potentially orchestrates fluid flow dynamics and renal oxygen supply. Recent work also demonstrated spontaneous regional synchronization of blood flow oscillations among over 100 neighboring star vessels across a large area of the rat renal cortex (71), which is regulated by TGF and abolished by furosemide (127). Accumulating evidence suggests that through TGF-mediated oscillations, a large number of nephrons can coordinate fluid flow via the vascular tree to achieve optimal control of blood flow over a large region (70) and sustain stable oscillations at the whole-organ scale. Considering that TGF and renin secretion share the same sensor at the MD (128, 129), TGF-mediated oscillations in the kidney can interact with the renin-angiotensin system to regulate systemic blood pressure (130). These findings, together with the work presented here, strongly suggest that further understanding of TGF-mediated oscillations in normal physiology, and how those oscillations are disrupted in renal pathophysiology, are critical to future clinical developments in the monitoring and treatment of acute kidney injury, chronic kidney disease, diabetic kidney disease, and a host of other conditions.

## Materials and Methods

### Sex as a biological variable

Both male and female animals are included in this study.

### Animal experiments

#### Intravital 2P Microscopy

All procedures were approved by the Institutional Animal Care and Use Committee of the National Cancer Institute and complied with the Guide for the Care and Use of Laboratory Animals. Male and female C57BL/6J mice (strain #000664) were purchased from Jackson Laboratories (USA) and bred in house. Animals were used at 1 month (N = 4), 3 months (N = 20) and 15 months (N = 4) of age. All mice were maintained in a pathogen-free and temperature-controlled facility, regulated with a 12-h light/dark cycle and fed *ad libitum* (standard chow diet; NIH-31 Open Formula, Envigo).

#### Resting-state MRI

All experiments were approved by the Institutional Animal Care and Use Committee at Washington University in St. Louis. Male (N = 6) and female (N = 6) Sprague-Dawley rats (10-11 weeks) were acquired from Charles River Laboratories (USA).

### *In vivo* surgery

Male and female mice were weighed and anesthetized with isoflurane (3%). After adequate anesthesia was ensured (confirmed by the absence of reflex in response to toe pinch), a 10-mm dorsal incision was made under sterile conditions, and the right kidney was exteriorized. The animal was placed on the stage of an inverted microscope with the exposed kidney placed on a coverslip-bottomed plate with a saline bath, and the kidney was visualized from below. During all procedures and imaging, anesthesia was administered (1.5-2% isoflurane) through a nosecone, and core body temperature was maintained in a heated chamber. Timelapse images of the renal cortex were acquired up to 150 mm deep below the surface. Water-soluble fluorophores were injected in a single bolus retro-orbitally to enable labeling of specific compartments in the living kidney: 70 kDa dextran-rhodamine B conjugate (2 mg/kg, Invitrogen, D1841) was used to label tubular fluid for hour-long tracking of tubular fluid dynamics. In some experiments, lucifer yellow was used (1 mg/kg, Invitrogen, L453) to label tubular fluid; 2M MW dextran-rhodamine B conjugate (1 mg/kg, Invitrogen, D7139) was used to label the circulating plasma. After the baseline imaging session, some animals received a second intravenous bolus of furosemide (1 mg/kg, Sigma-Aldrich, F4381) or empagliflozin (50 *μ*g/kg, MedChemExpress, HY-15409) or vehicle (0.2% DMSO in saline, Sigma-Aldrich, D2650), followed by a 30-minute imaging session to track the effect of pharmacological interventions on tubular fluid flow dynamics.

### Intravital two-photon (2P) microscopy

Timelapse images were acquired using a Leica SP8 two-photon fluorescence imaging system with a with a Leica 25X water-immersion objective (numerical aperture: 1.4) powered by a Chameleon Discovery laser at 760 nm (Coherent, Santa Clara, CA) and a DMI8 inverted microscope’s external Leica 4Tune spectral hybrid detectors (emission at 510-560 nm for lucifer yellow, 570-630 nm for Rhodamine B) (Leica Microsystems, Heidelberg, Germany). The potential toxicity of laser excitation and fluorescence to the cells was minimized by using a low laser power and high scan speeds to keep total laser exposure as minimal as possible. Timelapse images were collected as 2D time series (xyt, 2-5Hz framerate; 8-bit, 512×512 pixel) for 2-10 min with the Leica LAS X imaging software. The same instrument settings (laser power, offset, gain of both detector channels) were applied for imaging male and female mice.

### Intravital imaging data processing

Glomerular filtration of small to mid-sized fluorophores occurs is highly pulsatile *in vivo* (63, 65, 66), giving rise to periodic changes in fluorescence intensity at a given luminal region of interest (ROI). Relative changes in fluorescence intensity can be used as a proxy to infer SNGFR or tubular fluid flow dynamics. All timelapse images were processed with ImageJ. After the raw image file was imported, multiple luminal ROIs were selected manually and quantified for fluorescent intensity for all frames. Where minor motion artifacts were present, images were registered and corrected using the NanoJ-Core plugin (131, 132). In MATLAB (R2023b), the mean intensity of individual ROIs was subjected to Fast Fourier Transform (FFT) to determine dominant frequencies between 0.015 – 0.080 Hz (133). To remove secular shift and higher frequency components, detrending and smoothing was achieved by bandpass-filtering raw signals within the same frequency range. In some cases, reference sine waves of the dominant frequency were fitted to the raw signals for waveform check. Wavelet analysis was performed with the Python-based software pyBOAT (134).

### Immunofluorescence and confocal microscopy

Male and female mice were anesthetized with 250 mg/kg Xylazine, and 50 mg/kg Ketamine (diluted in saline) injected intraperitoneally (i.p.). Kidneys were fixed by transcardial perfusion of ice-cold PBS for 2 min followed by ice-cold 4% paraformaldehyde (PFA) in PBS at a rate of 5 ml/min. Kidneys were harvested and stored in 4% PFA in PBS overnight and processed in a sucrose gradient before embedding in OCT (Tissue-Tek). Blocks were kept at -80°C until 10*μ*m thick slices were sectioned with a cryostat. Slides were stored at -80°C until thawed, rehydrated, and blocked with 0.1% Triton X-100 and 10% FBS in PBS for 1 hr at room temperature. Slides were then incubated with primary antibodies at 4°C overnight. The following antibodies were employed in this study recognizing KIM-1 (Kidney Injury Molecule-1, Rat, 1:500, R&D Systems, MAB1817), FITC-conjugated LTL (Lotis Tetragonolobus Lectin, 1:1000, Vector Laboratories, FL-1321). In the following day, slides were washed three times for 15 minutes each, then incubated with secondary antibodies and Alexa Fluor 647 Phalloidin (1:1000, Invitrogen, A22287) for 1 hr at room temperature. After three 15-minute washes, sides were dried and mounted with Fluoromount-G and a coverslip. Tile scans and z-stacks of kidney sections were acquired using a Leica SP8 confocal laser scanning microscope using a 63x oil-immersion objective (numerical aperture: 1.4). Quantification of multi-channel fluorescence signals was performed with ImageJ using the default settings.

### Resting-state MRI

All imaging was performed on a Bruker 9.4 Tesla (T) small animal MRI system (Bruker, USA). Imaging and data processing were performed according to a published protocol (69). Briefly, animals were sedated throughout imaging using 1-2% isoflurane mixed with oxygen. Respiration, heart rate, and temperature were monitored and controlled to normal levels and to avoid aliasing of signals. Right kidney was imaged in all animals. For resting state imaging, a multi-slice 2D gradient recalled echo (GRE) - echo planar imaging (EPI) protocol was used with the follow parameters: echo time (TE)/repetition time (TR) = 7.55/150 ms, dummy scans = 107, voxel resolution = 0.5 x 0.5 x 0.5 mm^3^, and temporal resolution = 0.150 sec (6.67 Hz). Images were continuously acquired over 10 min. The whole kidney was manually segmented in images. Kidney cortex was defined as all voxels within 1.5 mm from the edge of the kidney. Power spectra were calculated for all voxel time series of MR signals using FFT. To compare with the mouse studies, a dominant frequency between 0.015 – 0.080 Hz was identified in the Fourier spectrum for each cortical voxel. In each animal, 250-300 cortical voxels were quantified and the mode (the most likely value) of the histogram of per-voxel dominant frequencies was chosen as the representative frequency.

### Renal blood flow data analysis

Original recordings in 6-month-old rodents from two studies (72, 78) were shared by the authors and processed locally in MATLAB (R2023b). In both studies, a transit-time ultrasound flow probe (Transonic Systems, United States) was mounted to the renal artery to measure renal blood flow in anaesthetized mice (72) and rats (78). We applied FFT to determine the dominant frequency between 0.015 – 0.080 Hz, and bandpass filtering was performed to visualize slow dynamics (< 0.1 Hz).

### Transdermal GFR measurement data analysis

Original recordings in 11-week-old and 30-week-old C57BL/6J mice from two studies (76, 79) were shared by the authors and processed locally in MATLAB (R2023b). The clearance profile after a single bolus of FITC-sinistrin is widely used for measuring real-time glomerular filtration rate using a transcutaneous sensor (MediBeacon GmbH, Germany) in conscious, free-moving animals. We applied FFT to determine the dominant frequency between 0.015 – 0.080 Hz.

### Statistical analysis

To include non-normally distributed datasets in our statistical analysis, we adopted non-parametric methods in this study. Specifically, resampling was used to determine statistical significance for between-group comparisons. In brief, data points from two groups were pooled and resampled 1 million times with replacement for large datasets. For small datasets (N < 10 in each group), we used a rank-based resampling method analog of the Mann-Whitney-Wilcoxon test. Resampled cases showing an absolute median difference larger than or equal to the original median difference were identified and enumerated. *P*-values were calculated by dividing the number of equal or more extreme cases by 1 million.

### Mathematical modeling

Historically, a number of mathematical models have been proposed to account for TGF-mediated oscillations in SNGFR and tubular fluid flow dynamics (83). These models vary widely in granularity and complexity. Harold Layton, Bruce Pitman and Leon Moore proposed a minimal mathematical model of TGF (84), consisting of a first-order partial differential equation describing sodium reabsorption along the TAL and an empirical negative feedback relation at the MD with an explicit time delay. Through mathematical analysis, their work provided fundamental insight into how oscillatory SNGFR can be sustained or abolished. However, to study male-female dynamic differences in TGF-mediated oscillations in SNGFR, it is necessary to track fluid flow and sodium handling in the PT. The tubular system can be represented as a single lumped compartment (135, 136) or as a one-dimensional form of the Navier-Stokes equations detailing individual constituents and complex interactions (137, 138). Here, we present a multi-compartment approach to model tubular hydrodynamics, striving to capture the underlying physical processes while keeping the system analytically and computationally tractable.

#### The compartmental model of TGF

Dividing the lumen of PT and TAL into arrays of smaller compartments (**Figure 3A**), we can track instantaneous fluid volume and sodium concentration in each compartment over time with the following statements: (1) SNGFR at the glomerulus determines the initial fluid flow rate into the first PT compartment; (2) Reabsorption of fluid in PT compartments is linear (first-order flow-dependent transport) and isotonic (fluid follows sodium transport across the epithelium, and [Na+] remains constant along PT compartments) (80–82); (3) Reabsorption of sodium occurs without fluid leaking through the epithelium in the TAL compartments. (4) [Na+] at the terminal TAL compartment is sensed at MD, eliciting negative feedback on SNGFR by controlling the vaso-elasticity of the afferent arteriole (AA).

Since fluid volume is conserved, changes in fluid volume in the *i*-th PT compartment (*V*_*i*_) are determined by the net effect of inflow, outflow and reabsorbed volume. The unidirectional convection along the tubule is modeled as cascading events of proportional fluid volume transfer specified by a rate constant *κ* (see schematic below). Thus, we have:

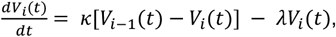

for *i* = 1, 2, …, *N*. Here, *κ* is a semi-phenomenological parameter inverse to the impedance to luminal fluid flow, which is controlled by the morphological features of the PT (such as the lumen radius) as well as by the fluid composition (such as solute density and viscosity); *λ* represents the PT sodium reabsorption rate, which is controlled by net sodium transport efficiency and cellular metabolic capacity to support transport processes.

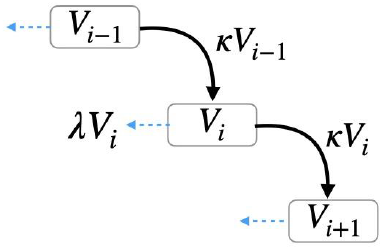

Considering that the total amount of sodium is conserved, changes in [Na+] in the *j*-th TAL compartment (*C*_*j*_) are determined by the net change of inflow, outflow and reabsorption of sodium (84):

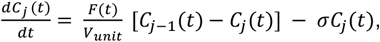

for *j* = 1, 2, …, *M*; where *F*(*t*) = *κV*_*N*_(*t*), *C*_1_(*t*) = *c*_*i*_. Here, *σ* represents the TAL sodium reabsorption rate; *c*_*i*_ refers to the [Na+] in the plasma.

Lastly, changes in fluid volume feeding into the first PT compartment are governed by the negative feedback from MD to AA, modeled as a downward-going sigmoidal function:

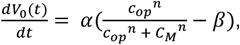

where SNGFR = *kV*_0_(*t*) and *C*_*M*_ refers to [Na+] in the last TAL compartment. Here, *c*_*op*_ represents the operating [Na+] at MD, which can be seen as the optimal [Na+] for which no effect of vaso-constriction nor vaso-dialation would be elicited; *n* refers to the maximal slope of the sigmoidal function, which is controlled by the sensitivity of the negative feedback from MD to AA; *α* is the scaling factor for the sigmoidal function, and *β* is the vertical offset.

#### Numerical simulation and analysis

The compartmental model of TGF was parameterized according to **Table S1** in the *Supplementary Materials*, which corresponds to a typical male mouse nephron producing a self-sustained oscillation in SNGFR at 0.04 Hz (average SNGFR = 4.27 *nl/min*). Unit volume (0.005 *nl*) was chosen for a typical TAL compartment with a lumen radius of 10 *μ*m and a width of 16 *μ*m (about the length spanning 3 tubular epithelial cells), and we used a time step of 0.1 sec during simulation.

Simulations were performed in MATLAB (R2023b) using the `ode15s` function. The compartmental model of TGF was simulated for 10 minutes (600 seconds), for different parameter combinations of *λ* and *κ*, while keeping other parameters constant. System behaviors over the last 200 seconds were used to calculate the average SNGFR, amplitude and frequency of oscillatory SNGFR if any, as well as the fractional reabsorption of sodium in the PT.

## Supporting information

TGF_SexDiff_MS_Supp

## Additional Information

## Acknowledgements

We thank all members of the Porat-Shliom laboratory for helpful comments on experimental design for intravital two-photon microscopy. We are very grateful that Drs. William A. Cupples and Keisa W. Mathis shared renal blood flow datasets with us, and that Drs. Eryn E. Dixon and Jorge F. Giani shared recordings from transdermal GFR measurements with us. We also thank Drs. Donald J. Marsh, Janos Peti-Peterdi, Scott C. Thomson, Mark A. Knepper, Christopher S. Wilcox, Petter Bjornstad and Adriana C. C. Girardi for stimulating discussions.

## Author Contributions

Conceptualization: LX, AG, NPS, EJD

Methodology: LX, AG, KMB, EJB, KB, AE, AAM, NPS, EJD

Formal Analysis: LX, EJB

Investigation: LX, EJB, LB

Data curation: LX

Software: LX

Visualization: LX, EJB

Project administration: LX

Resources: LX, AG, KMB, VV, AE, AAM, NPS, EJD

Supervision: AG, KMB, NPS, EJD

Writing – original draft: LX

Writing – review and editing: LX, AG, KMB, VV, AE, AAM, NPS, EJD

Funding acquisition: AG, KMB, NPS, EJD

## Data and Code Availability

All data needed to evaluate the conclusions in the paper are present in the paper and the Supplementary Materials. All original codes and simulation results are available on GitHub: https://github.com/LingyunXiong/TGF_SexDiff

## Supplementary Materials

### Supplementary Figures

Figure S1. Detecting physiological oscillations across spatial scales in the kidney (related to Figure 1).

Figure S2. Male-female frequency differences in TGF-mediated oscillations (related to Figure 2).

Figure S3. A compartmental model of TGF recapitulates essential nephron physiology (related to Figure 3).

Figure S4. Loss of TGF-mediated oscillations underlies male-heightened risk (related to Figure 4).

### Supplementary Tables

Table S1. Initial conditions and parameter values for computational simulations

Table S2. Commercial reagents used in this study

