## Supplementary material for "Life in the fast lane: Functional consequences of male-female dynamic differences in the renal auto-regulation of flow": TGF_SexDiff_MS_Supp

Lingyun Xiong *et al.*

**This PDF file includes:**

Figures S1 to S4

Tables S1 to S2

**Figure S1. Detecting physiological oscillations across spatial scales in the kidney**

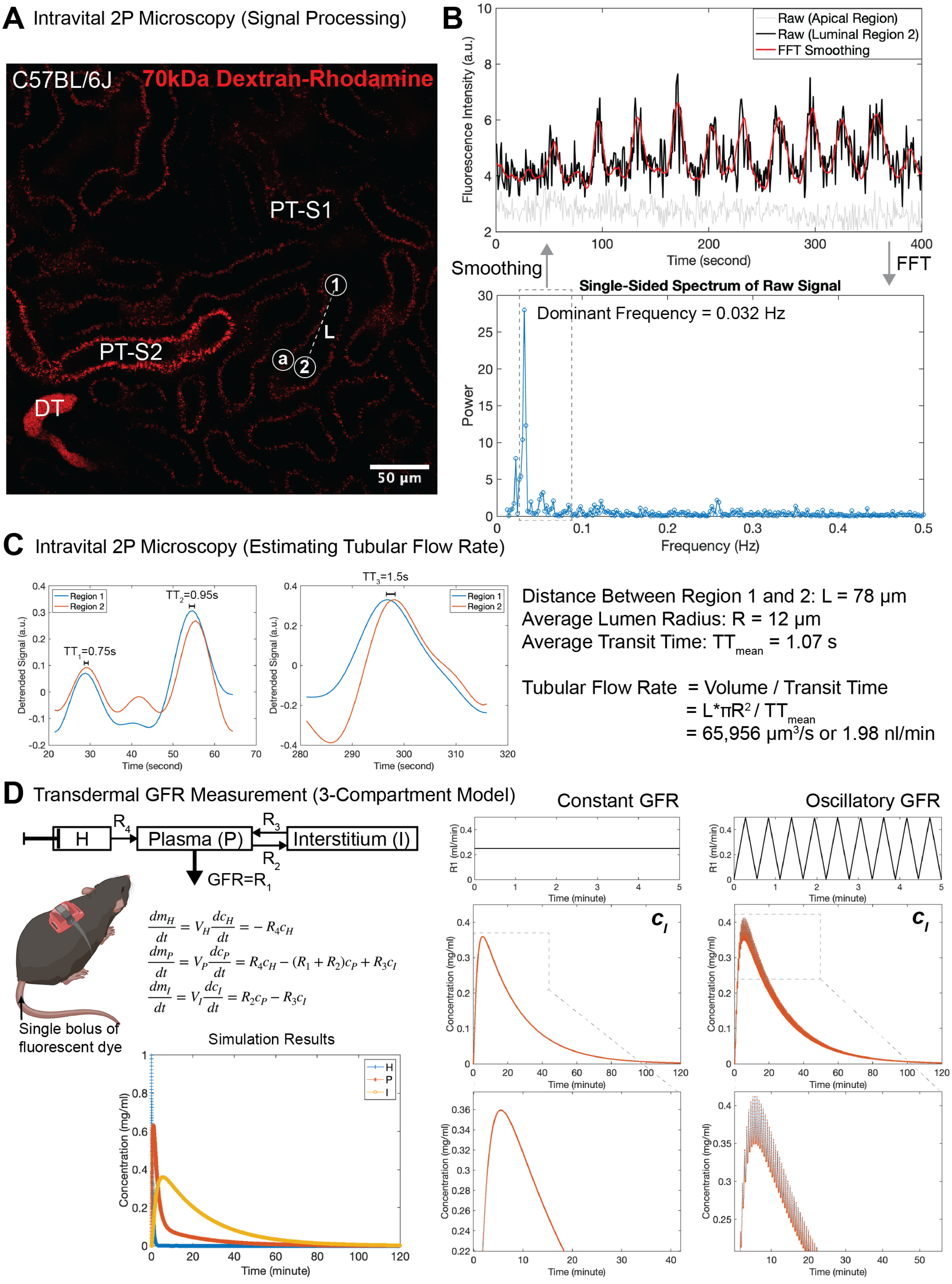


$$\frac{dm_H}{dt} = V_H\frac{dc_H}{dt} = -R_4c_H$$
$$\frac{dm_P}{dt} = V_P\frac{dc_P}{dt} = R_4c_H - (R_1+R_2)c_P + R_3c_I$$
$$\frac{dm_I}{dt} = V_I\frac{dc_I}{dt} = R_2c_P - R_3c_I$$

**Figure S1. Detecting physiological oscillations across spatial scales in the kidney (related to Figure 1).** (A) A representative field of view of the kidney cortex of an anaesthetized male C57B/6J mouse injected with a bolus of 70kDa dextran-rhodamine, where two luminal ROIs and one apical ROI are marked. (B) Workflow of signal processing for intravital two-photon microscopy. Top: Recording of a spontaneous oscillation in the intensity of 70kDa dextran-rhodamine within a luminal ROI in the kidney of an anaesthetized male C57B/6J mouse. Bottom: Fast Fourier transform (FFT) was used to determine the dominant frequency in the raw signal. Detrending and smoothing was achieved by bandpass-filtering raw signals within a frequency range of choice. (C) Estimating tubular flow rate from intravital two-photon microscopy. Detrended signals in the two ROIs (as in A), where transit time can be estimated by delays in signal peaking times. Based on distant measurements between the two ROIs and average lumen radius (92), average tubular flow rate is calculated by the fluid volume / transit time (65). (D) Modeling transdermal GFR measurement by simulating the 3-compartment model (77) indicated that compared to constant GFR input, oscillatory GFR input results in oscillatory fluorescence intensity in the interstitial compartment, which can give rise to the oscillatory clearance profile shown in Figure 1E.

**Figure S2. Male-female frequency differences in TGF-mediated oscillations**

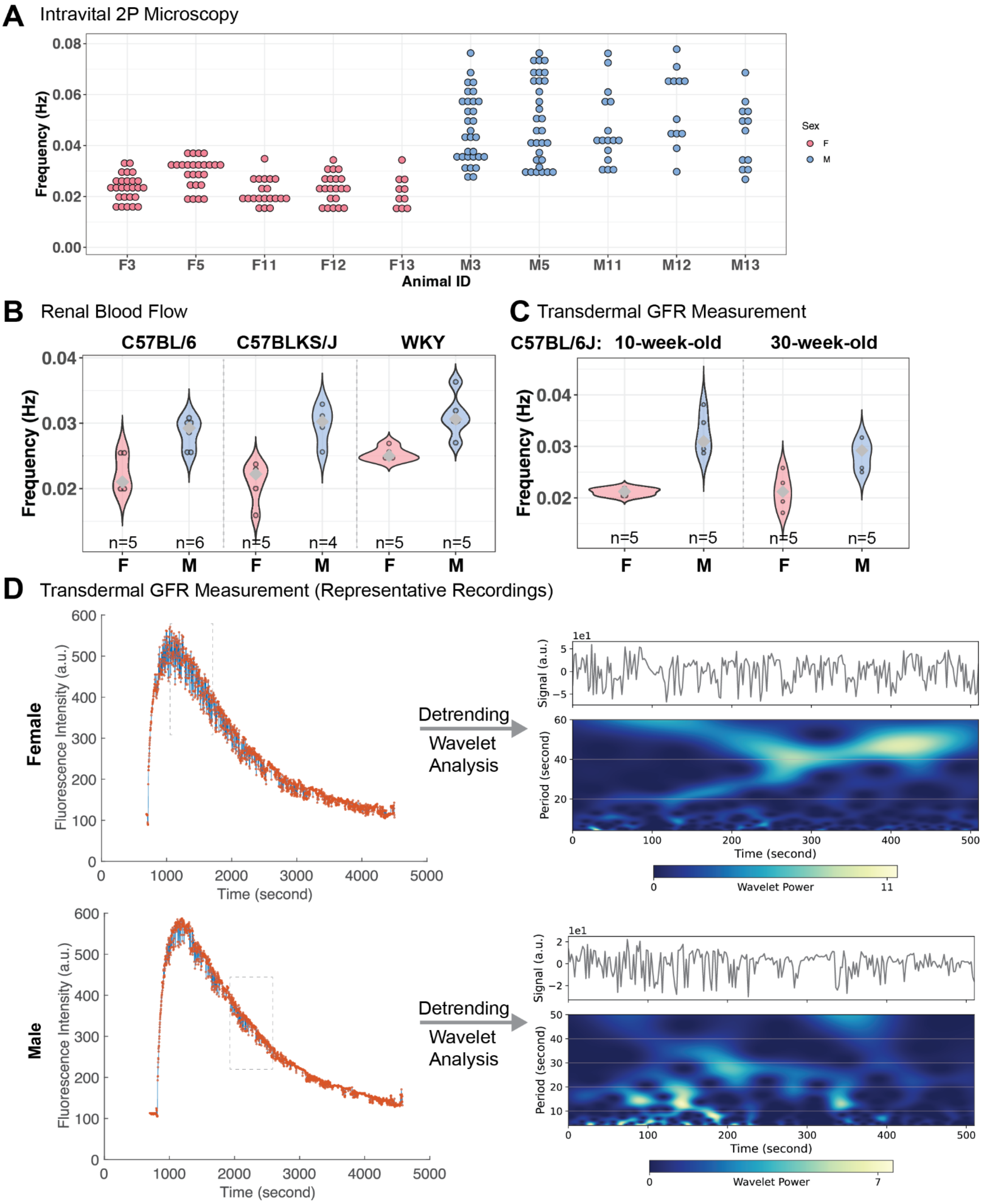


**Figure S2. Male-female frequency differences in TGF-mediated oscillations (related to Figure 2).** (A) Distribution of single-nephron dominant frequency in individual 3-month-old male and female C57BL/6J mice, measured by intravital two-photon microscopy. Multiple regions were quantified in each animal and all values are shown. (B) Distribution of whole-kidney dominant frequency in 6-month-old male and female rodents (C57BL/6 & C57BLKS/J mice and WKY rats), quantified from transit-time ultrasound-based renal blood flow data (72, 78). Sample number indicates total numbers of animals used. (C) Distribution of whole-body clearance dominant frequency in C57BL/6J mice from two age groups: 10-week-old (76) and 30-week-old (79). (D) Wavelet analysis was used to extract time-dependent periodicity in transdermal GFR measurement from 10-week-old male and female C57B/6J mice.

**Figure S3. A compartmental model of TGF recapitulates essential nephron physiology**

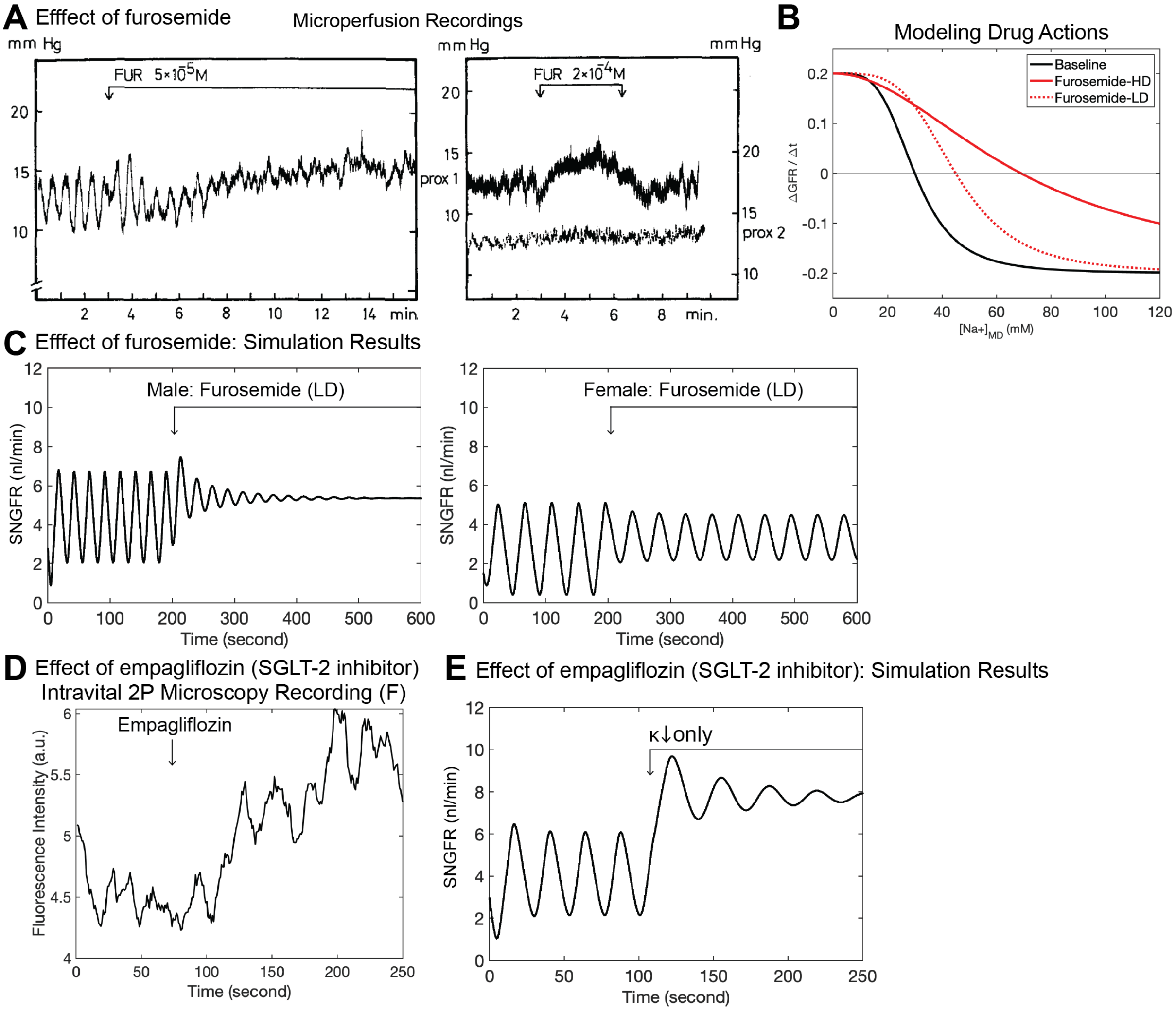


**Figure S3. A compartmental model of TGF recapitulates essential nephron physiology (related to Figure 3).** (A) The effect of furosemide at high and low doses on proximal tubular pressure (reprinted from ref. (86), with permission). (B) The dose-dependent effect of furosemide can be modeled by modifying the sigmoid function with different levels of desensitization at MD. (C) Simulation results of low-dose furosemide for typical male and female $\lambda - \kappa$ parameter combinations. (D) The acute effect of empagliflozin on luminal fluorescence intensity in a 3-month-old female C57BL/6J mouse. (E) Simulating the acute effect of empagliflozin by decreasing $\kappa$ alone was insufficient to reproduce the experimental observation shown in Figure 3E.

**Figure S4. Loss of TGF-mediated oscillations underlies male-heightened risk**

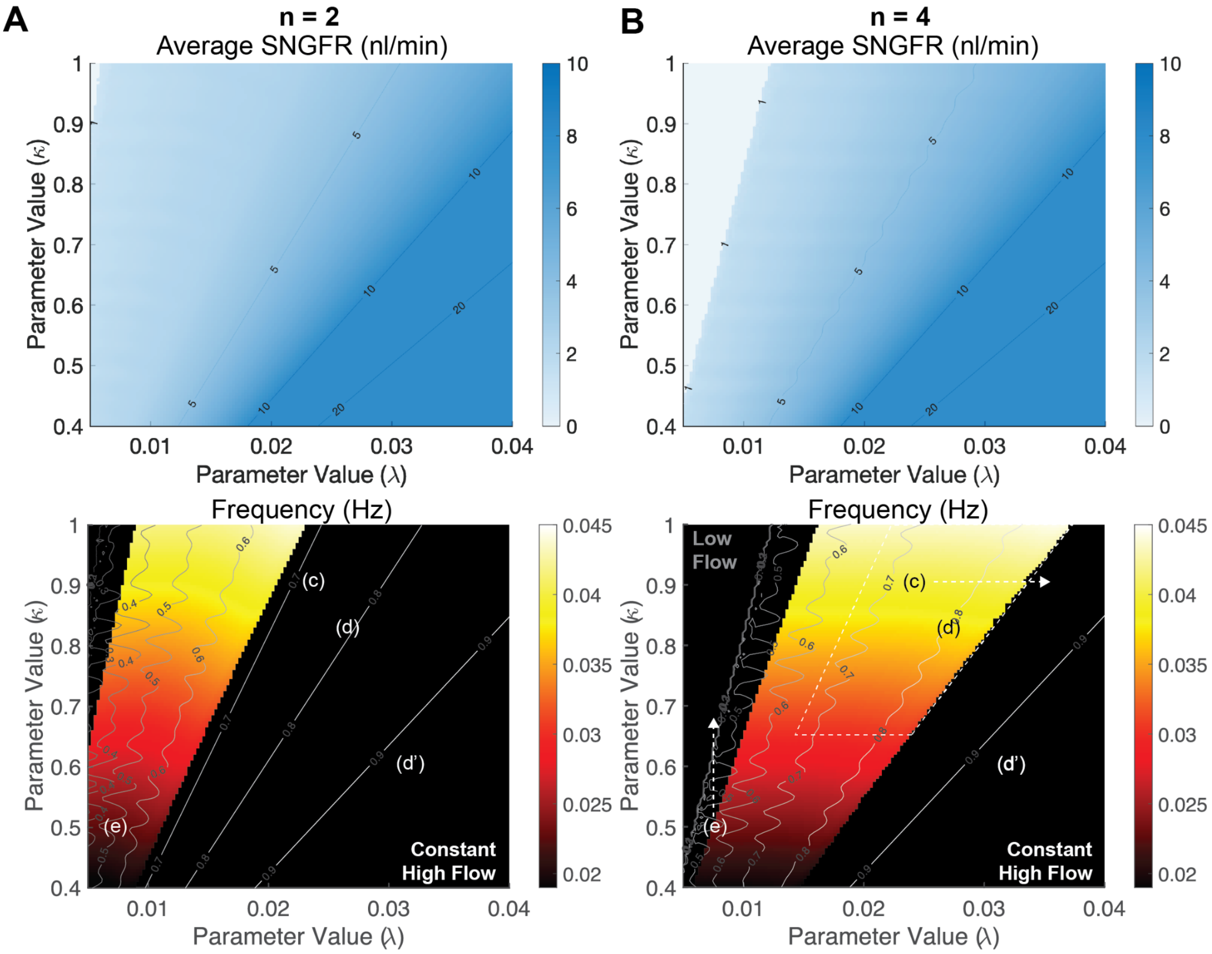


**Figure S4. Loss of TGF-mediated oscillations underlies male-heightened risk (related to Figure 4).** Numerical simulations were performed for different combinations of $\lambda$ and $\kappa$, while keeping a 'shallower' slope ($n$=2; A) or a 'steeper' slope ($n$=4; B) of the sigmoid function at MD. Heatmap on top shows the distribution of average SNGFR across the 2D parameter space. Heatmap at the bottom indicates the frequency of oscillatory behavior, overlaid with the contour plot indicating PT fractional reabsorption of sodium.

**Table S1. Initial conditions and parameter values for computational simulations.**

| Variable | Description | Initial Value |
|---|---|---|
| $V_i(t)$ | Fluid volume in the $i$-th PT compartment | $0.05\ nl$ |
| $C_j(t)$ | [Na+] in the $j$-th TAL compartment | $150\ mM$ |
| **Parameter** | **Description** | **Value** |
| $c_i$ | [Na+] in the plasma | $150\ mM$ |
| $c_{op}$ | Operational [Na+] at the MD | $30\ mM$ |
| $\alpha$ | Scaling factor for the sigmoidal function | $0.02\ nl\ s^{-1}$ |
| $\beta$ | Vertical shift of the sigmoidal function | $0.5$ |
| $n$ | Maximal slope of the sigmoidal function | $3$ |
| $\kappa$ | PT fluid flow rate constant | $9.2\ s^{-1}$ |
| $\lambda$ | PT sodium reabsorption rate | $0.24\ s^{-1}$ |
| $\sigma$ | TAL sodium reabsorption rate | $1\ s^{-1}$ |
| $N$ | Number of PT compartments | $50$ |
| $M$ | Number of TAL compartments | $60$ |

**Table S2. Commercial reagents used in this study**

| REAGENT | SOURCE | IDENTIFIER |
|---|---|---|
| **Antibodies** | | |
| FITC-conjugated Lotis Tetragonolobus Lectin (LTL) | Vector Laboratories | Cat# FL-1321; RRID: AB_2336559 |
| Rat monoclonal anti-KIM1 | R&D Systems | Cat# MAB1817; RRID: AB_2116445 |
| **Fluorescent Probes** | | |
| Alexa Fluor 647 Phalloidin | Invitrogen | Cat# A22287 |
| 70-kDa dextran-rhodamine B conjugate | Invitrogen | Cat# D1841 |
| 2M MW Dextran, Tetramethyl-rhodamine | Invitrogen | Cat# D7139 |
| Hoechst 33342 | Invitrogen | Cat# H1399 |
| Lucifer Yellow CH | Invitrogen | Cat# L453 |
| **Pharmacological Reagents** | | |
| DMSO | Sigma-Aldrich | Cat# D2650 |
| Furosemide | Sigma-Aldrich | Cat# F4381 |
| Empagliflozin | MedChemExpress | Cat# HY-15409 |
